# Embryonic organizer specification in the mud snail *Ilyanassa obsoleta* depends on intercellular signaling

**DOI:** 10.1101/2023.05.23.541988

**Authors:** Jessica E. Wandelt, Ayaki Nakamoto, Morgan Q Goulding, Lisa M. Nagy

## Abstract

In early embryos of the neogastropod *Ilyanassa obsoleta*, cytoplasmic segregation of a polar lobe is required for establishment of the D macromere, empowering its great-granddaughter macromere 3D to act as a single-celled organizer that induces body patterning along the secondary axis. We present evidence that polar lobe inheritance is not sufficient to specify 3D potential, but rather makes the D macromere lineage responsive to some intercellular signal(s) required for activating the special properties of 3D. Experimental removal of micromeres results in loss of organizer-linked MAPK activation, complete and specific defects of organizer-dependent larval organs, and progressive cell cycle retardation leading to equalization of the normally accelerated division schedule of 3D. Ablation of the second-quartet micromere 2d greatly potentiates the effects of first-quartet micromere ablation. Our findings link 3D establishment in *I. obsoleta* to the putative ancestral spiralian mechanism in which a signal from micromeres leads to specification of 3D among four initially equivalent macromeres.

**Summary Statement:** Cell ablation experiments on embryos of the snail *Ilyanassa* reveal that specification of the organizer cell 3D depends on cell-extrinsic cues as well as polar lobe inheritance.

## INTRODUCTION

In spiralian development, a deeply conserved cleavage pattern is accompanied by a conserved pattern of cell lineage specification. The first two mitotic divisions produce four body-quadrant founder cells (A, B, C, D), which then synchronously undergo three rounds of asymmetric division, forming three quartets of micromeres that collectively give rise to the entire ectoderm. Intrinsic difference in potential between the three micromere quartets creates a pattern along the primary (animal-vegetal) embryonic axis. Overlaid on this pattern is a gradient of intercellular signaling along an orthogonal secondary axis, initiated by one ‘organizer’ cell of the D quadrant. This two-step body-patterning mechanism was revealed by experiments on distantly related gastropod mollusks (Clement, 1976; van den Biggelaar, 1977; Arnolds et al., 1983; Lambert, 2010): in every snail taxon examined, organizer activity begins in the 3D macromere (or its daughter 4d) (Clement, 1962; Labordus and van der Wal, 1986; Martindale, 1986; Henry et al., 2006), and is linked to activation of a mitogen-activated protein kinase (MAPK) exclusively in the 3D cell (Lambert and Nagy, 2001; Lambert and Nagy, 2003; Koop et al., 2007; Henry and Perry, 2008; Tan el al., 2022).

In most of these snails, the first two rounds of mitosis produce four body-quadrant founder cells which are equipotent. After formation of the three micromere quartets, four third-order macromeres lose equipotency when one of them adopts the identity of 3D, activates MAPK and initiates organizer signaling. The crucial selection of just one macromere as 3D depends on signaling from head precursor cells (first-quartet micromere derivatives) surrounding the animal pole. Ablation of one or two selected first-quartet cells imposes a strong spatial bias on 3D selection among the macromeres (van den Biggelaar and Guerrier, 1979; Arnolds et al., 1983); further, 3D specification can be entirely blocked by killing the whole first quartet (van den Biggelaar and Guerrier, 1979; Boring, 1989; Freeman and Lundelius, 1992; Henry et al., 2006). The inductive action of micromeres in 3D specification also occurs in basal mollusks (van den Biggelaar, 1996) and in other phyla among Lophotrochozoa (Boyer, 1989; Henry, 2002; Edsinger-Gonzales et al., 2007).

In Neogastropoda (a very large clade including whelks, winkles, and cone snails), a novel mechanism of 3D specification appears to have evolved (Freeman and Lundelius, 1992). During the first two divisions, a transiently extruded ‘polar lobe’ (PL) shunts material from the egg’s vegetal region to one daughter cell; seminal experiments on the neogastropod *Ilyanassa obsoleta* indicated that PL inheritance confers D-lineage identity on one quadrant-founder cell (Crampton, 1896; Clement, 1952). This apparently obviates the ancestral 3D-inducing signal: ablation of all four first-quartet micromeres in *I. obsoleta* was reported to yield headless but otherwise perfect larvae (Sweet, 1998), indicating undisturbed organizer function and implying cell-autonomous specification of 3D.

We set out to test the influence of micromeres on 3D specification in *I. obsoleta* more stringently. Using four independent assays of 3D identity or organizer function, we assessed effects of multiple cell ablations in two contexts: intact embryos and quarter-embryo fragments. We found that 3D specification in *I. obsoleta* requires intercellular signaling as well as PL inheritance. The 3D-activating signal appears to arise from a combination of multiple first-quartet cells and at least one second-quartet cell (2d).

## RESULTS AND DISCUSSION

### Polar lobe inheritance is required for MAPK activation in a third-order macromere

In early embryos of *I. obsoleta*, reactivity to an antibody against the kinase-active form of MAPK (ERK1/2) first appears in the 3D macromere shortly after its birth and persists until the cell divides about ninety minutes later; no other cells express this immunoreactivity prior to 3D. Pharmacological interference with the earliest period of MAPK activation blocks ectodermal organogenesis that is normally induced by the 3D organizer (Lambert and Nagy, 2001). We therefore focused on expression of active MAPK in 3D as a proxy for activation of organizer function. First, we tested the assumption that 3D MAPK activation depends on polar lobe inheritance by the D cell at second cleavage. When quadrant founder cells (A, B, C, or D) were cultured after isolation at the four-cell stage and immunostained for active MAPK about eighty minutes after formation of their third micromeres, only the 3D cell showed a positive result, as expected (Fig. 1A-C). We also observed absence of MAPK activation in quarter-embryos isolated after PL removal at first cleavage. Among 71 non-PL-inheriting quadrant isolates, none showed MAPK activation.

**Figure 1.**
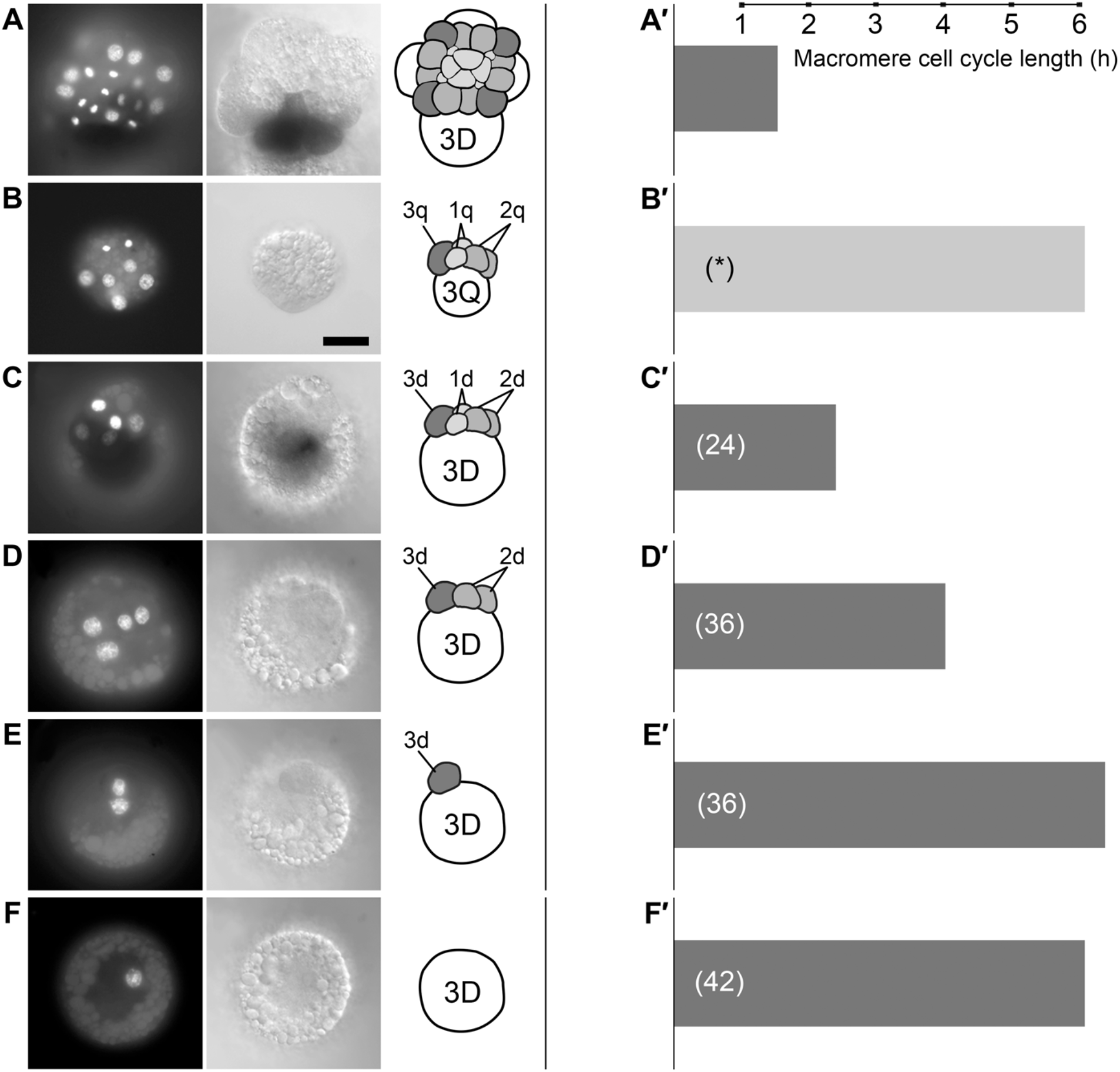
Micromere ablation effects on 3D MAPK activation and cell cycle timing in quadrant isolates. (A-F) Sibling whole and partial embryos assayed for activated MAPK at 3q+80 minutes. Left: DAPI-stained representative embryos. Middle: DIC images of same embryos showing activated MAPK detected with HRP. Right: cartoons showing cells present. (A) Activated MAPK in intact control embryo. (B) Absence of activated MAPK in unidentified quadrant (‘Q’) isolated after polar lobe removal. (C) MAPK activation in D quadrant isolate. (D) Absence of activated MAPK in D quadrant isolate lacking 1d. (E) Absence of activated MAPK in D quadrant isolate lacking 1d and 2d. (F) Absence of activated MAPK in D quadrant isolate lacking 1d, 2d, and 3d. (A#-F#) Bars representing average cell cycle length (time to fourth micromere formation) in classes of control and operated embryos shown in (A-F). Numbers (#) of embryos are indicated. Asterisk (*) in B# signifies that 3Q division timing in non-PL-inheriting quadrant isolates was not measured in this study; the value shown represents cell cycle duration in PL-deleted embryos (Clement, 1952). Scale bar in B, 50 μm.

### MAPK activation in isolated D quadrants depends on presence of micromeres

Aiming to identify possible extracellular influence on 3D organizer function, we examined MAPK activation in an idealized experimental setup with a minimum of cells present. D quadrant founder cells isolated at the four-cell stage proceed through cleavage and subsequent development with only minor deviation from *in situ* behavior (Crampton, 1896; Clement, 1956; Goulding, 2003). To assess the autonomy of 3D organizer function, we repeated the isolation of D quadrant founder cells, then additionally removed one or more micromeres (1d/2d/3d), subsequently assaying for activation of MAPK in 3D (Fig. 1D-F). In the great majority of whole D quadrant isolates (106/115), MAPK was activated as normal in the 3D macromere. Following ablation of the 1d micromere, however, only 29/113 D quadrant isolates showed MAPK activation in 3D (Fig. 1D). Successive removal of both first and second quartet micromeres (n=39), or all three micromeres (n=22) completely prevented 3D MAPK activation (Fig. 1E, F).

### Normal precocity of 3D mitosis depends on presence of micromeres

Another conserved aspect of 3D behavior is a cell division pattern distinct from its cousins in the A, B, and C quadrants. Following MAPK activation, 3D uniquely divides to produce the mesentoblast 4d (founder of most of the mesoderm, as well as the hindgut). In *I. obsoleta*, this division occurs three to four hours before the synchronous division of 3A, 3B, and 3C (Clement, 1952, Goulding, 2009). PL removal abolishes this difference, causing all 3Q cells to divide with the long delay of 3ABC (and yielding all non-mesentoblast daughters) (Clement, 1952). While examining D quadrant isolates for MAPK activation, we noticed that 3D division was retarded in every class of partial embryo. We therefore examined division timing closely in another set of experiments. In D isolates cultured intact with all D quadrant micromeres (n=24), 3D divided approximately 50 minutes later than 3D in whole embryos from the same brood (Fig.1A′-C′). In D isolates lacking the 1d cell (n=36), the abnormal 3D cell cycle delay was prolonged to almost 150 minutes (Fig. 1D′). Following ablation of both 1d and 2d (n=36) 3D division was delayed by almost 300 minutes (Fig. 1E′): in this situation 4d formation coincided with the timing of 4abc formation in whole embryos. The same long delay of 3D division resulted after ablation of all three micromeres (n=42) (Fig. 1F′).

We next examined 3D division timing after removal of selected micromeres from whole embryos (Fig. 2A-F). Ablation of either a single micromere 1d (n=20) or 2d (n=19), or the combination of 1d and 2d (n=60) had little effect on 3D division timing: in all cases 3D divided with 20 minutes of controls. By contrast, ablation of the entire first quartet (1a, 1b, 1c, and 1d; collectively abbreviated ‘1q’) resulted in prolongation of the 3D cell cycle by approximately 70 minutes (n=88). Strikingly, ablation of 1q and then 2d resulted in 3D division delay of nearly five hours (n=29), coinciding with division of 3ABC macromeres in the same embryos as well as unoperated controls.

**Figure 2.**
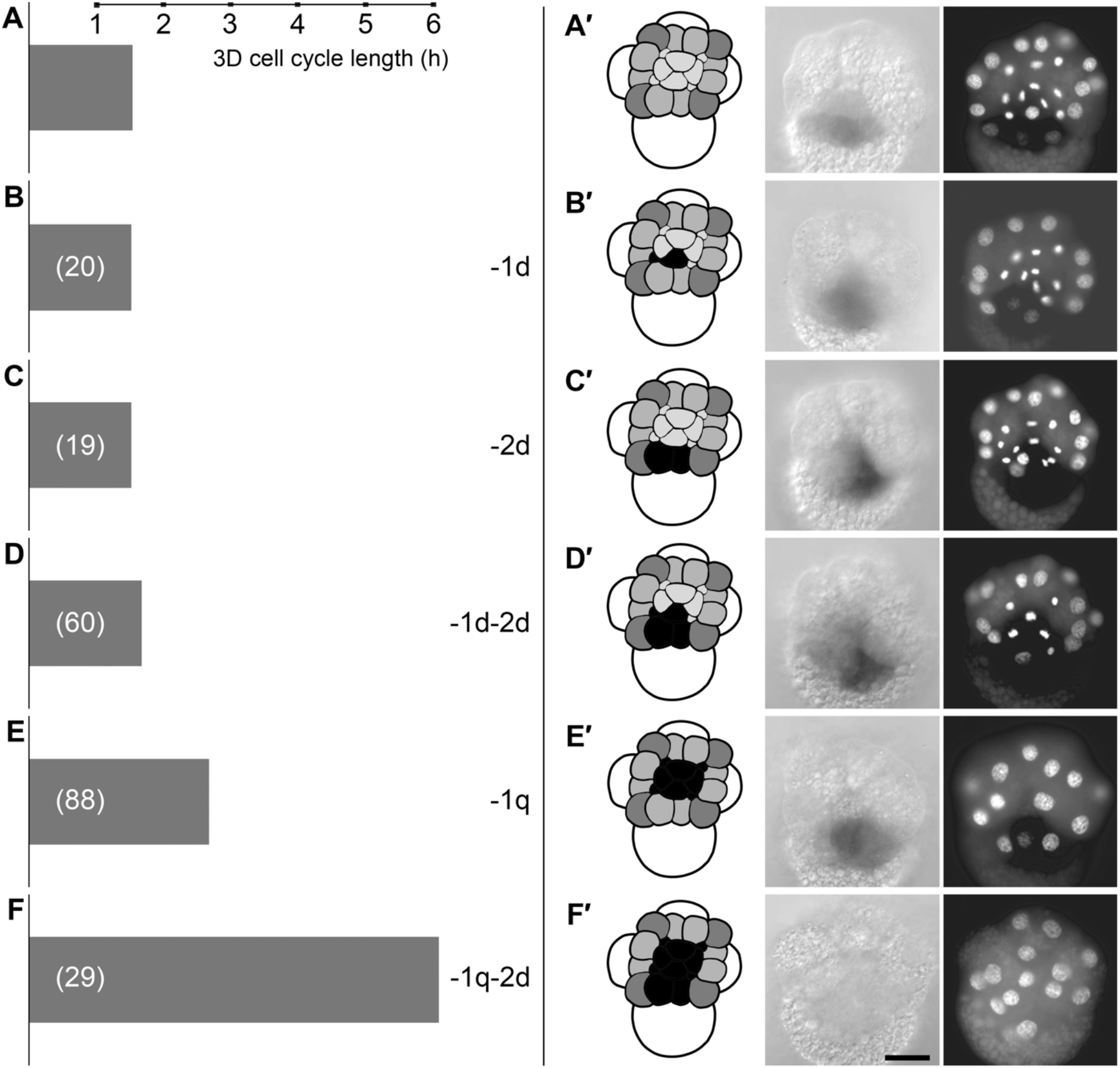
3D cell cycle timing and MAPK activation depend on multiple micromeres. (A-F) Bars representing average 3D cell cycle length in classes of control and operated embryos shown in (A′-F′). (A′-F′) Co-cultured sibling whole and partial embryos assayed for activated MAPK at 3q+80 minutes. Left: DAPI-stained representative embryos. Middle: DIC images of same embryos showing activated MAPK detected with HRP. Right: cartoons showing cells present, with ablated micromeres colored black. (A#) Control embryo. (B#) Embryo lacking 1d. (C#) Embryo lacking 2d. (D#) Embryo lacking 1d and 2d. (E#) Embryo lacking entire first quartet (‘1q’). (F#) Embryo lacking 1q and 2d. Scale bar, 50 μm.

### 3D MAPK activation depends on presence of both first- and second-quartet micromeres

Given the different effects observed after ablation of 1d and 2d in whole embryos compared to the same double ablation in D quadrant isolates, we extended the assay of 3D MAPK activation by removing single or multiple micromeres from whole embryos. Ablation of 1d (n=32) or 2d (n=42) had no apparent effect on 3D MAPK activation (Fig. 2A′-F′). Successive ablation of both 1d and 2d resulted in detectable 3D MAPK activation in only 74% of embryos (n=85). Following ablation of the entire first quartet (1q), 3D MAPK activation was detectable in 59% of embryos (n=58). Again, ablation of both 1q and 2d yielded a much stronger effect, resulting in absence of 3D MAPK activation in all embryos examined (n=26).

### 3D-induced ectodermal MAPK activation depends extrinsically on 1q and 2d

Shortly following 3D MAPK activation, *I. obsoleta* embryos exhibit a second wave of 3D-dependent MAPK activation in ectodermal precursors that are in contact with 3D; these cells’ 3D-dependent fates are also sensitive to an inhibitor pulse during the time interval of their own MAPK activation (Lambert and Nagy, 2001). We therefore asked if ectodermal MAPK activation is sensitive to early removal of 1q and 2d micromeres. Following ablation of just 1d (n=16) or 2d (n=18), all embryos showed activated MAPK in the remaining dorsal and lateral micromeres (Fig. 3A-C). The same was seen in all embryos lacking both 1d and 2d (n=17) (Fig. 3D) or the entire first quartet (n=32) (‘1q’ in Fig. 3E). Once more, the effect of removing both 1q and 2d was striking, with detectable micromere MAPK activation abolished in 20/21 embryos (Fig. 3F).

**Figure 3.**
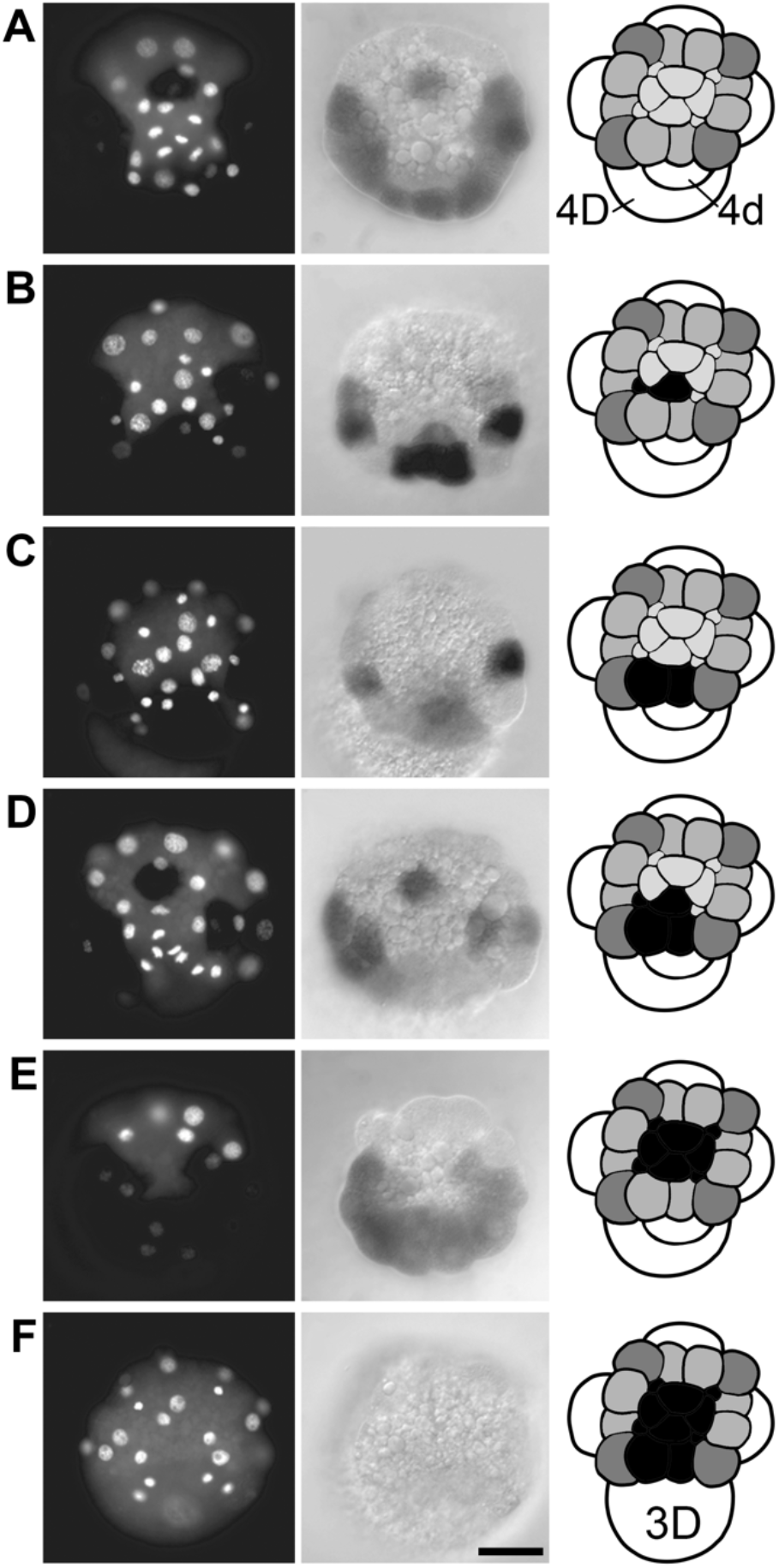
Ectodermal MAPK activation after early micromere ablations. Sibling whole and partial embryos assayed for activated MAPK at 3q+210 minutes. (A) Control embryo. (B) Embryo after 1d ablation. (C) Embryo after 2d ablation. (D) Embryo after ablation of 1d and 2d. (E) Embryo after ablation of entire first quartet (1q). (F) Embryo after ablation of 1q and 2d. Cartoons show ablated cells colored black. Note presence of 4d cell in (A-E). Scale bar, 50 μm.

### Organizer-dependent ectodermal organogenesis depends extrinsically on 1q and 2d

Extending the assessment of organizer function, we followed ablation of first- and second-quartet cells by culturing to near-hatching larva stage, assaying development of organizer-dependent trunk ectoderm (shell and foot) (Atkinson, 1971). Phenotypes are exemplified in Figure 4 (A-D). After first-quartet (1q) ablation, a third of all operated embryos (21/65) developed a flawless trunk and a complete absence of head structures (i.e., normally 1q-derived tissue), as previously reported (Sweet, 1998). We refer to this ‘headless’ phenotype, which indicates full functioning of the 3D organizer, as Type I. In another 15% of 1q-ablated embryos (Type II), shell and foot structures developed incompletely and with deranged spatial organization. Strikingly, most 1q-ablated embryos (52%) formed neither shell/foot structures nor a recognizable body axis; this phenotype (Type III) indicates complete loss of 3D organizer function. Hence, organizer function is perturbed by 1q ablation. Yet again, the additional ablation of 2d severely worsens the body-patterning performance, resulting almost exclusively in Type III development (Fig. 4E).

**Figure 4.**
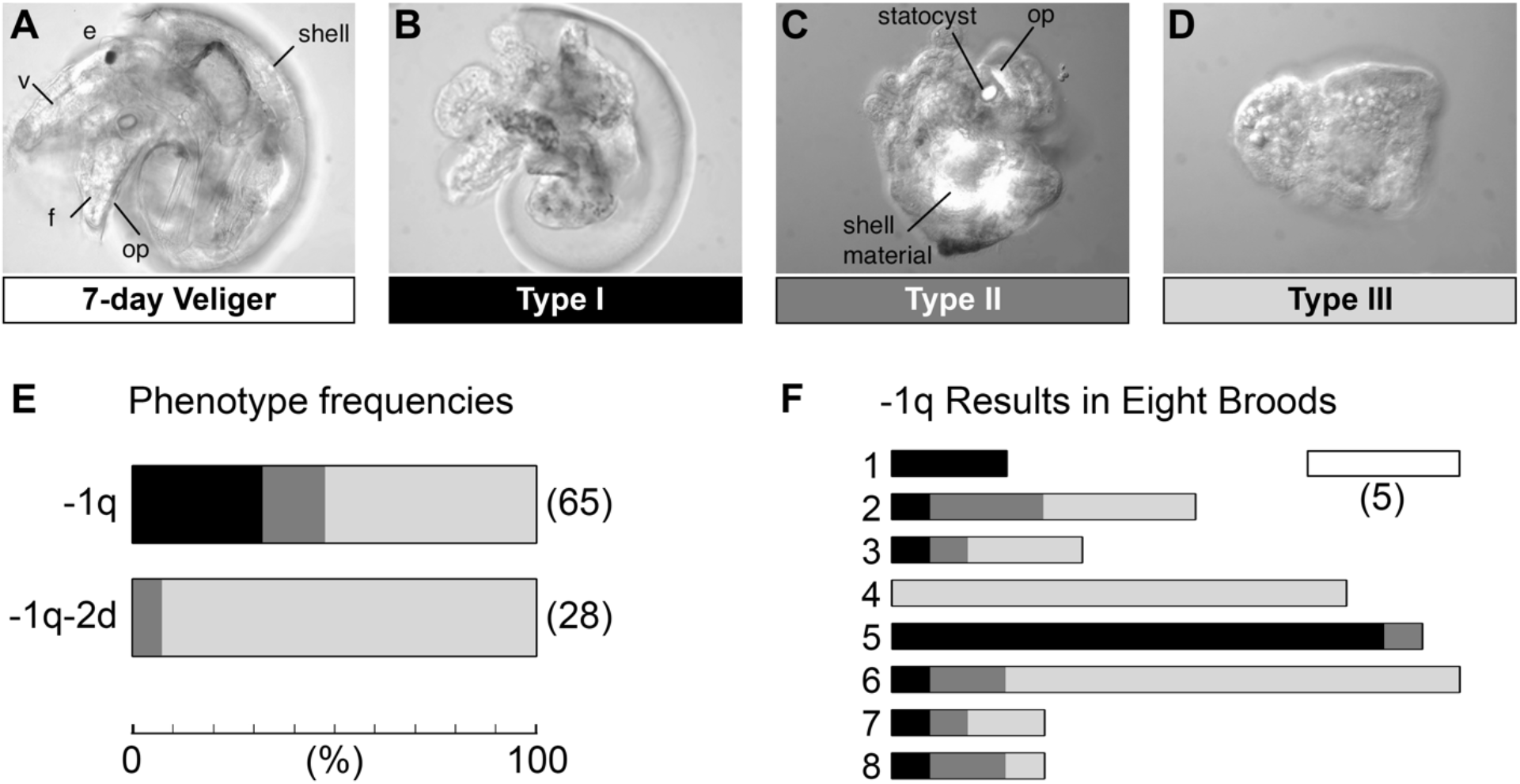
Multiple-micromere ablation disrupts organizer-induced organogenesis, with effects varying between broods. (A-D) Representative images of experimental embryos scored for larval morphogenesis and differentiation at seven days of development. (A) Unoperated control; near-hatching-stage veliger. Left side view, anterior to the left, showing one of the eyes (e), velum (v), foot (f) with operculum (op), and shell. (B) Type I. Left side view, anterior to the left. Trunk structures appear normal; head structures are absent. (C) Type II. No evident dorsoventral axis. Several tissue structures (internal shell mass, partial operculum, statocyst) are revealed by birefringence under polarized light. (D) Type III. No dorsoventral axis, no evident organs. (E) Phenotype frequencies after ablation of entire first quartet (−1q) or sequential ablation of 1q and 2d (−1q-2d). Total numbers (#) of embryos are indicated. (F) Numbers of 1q-ablated embryos showing each phenotype in eight broods. Phenotypes are differentially shaded: Type 1 is black, Type II is dark gray, Type II is pale gray. Scale bar, five embryos.

### Large interbrood variation in effect of 1q ablation

Our results disagree with those reported in a previous study of *I. obsoleta*, where 1q ablation yielded only headless larvae with well-formed trunks (Sweet, 1998). In our experiments, 1q ablations were done on embryos from eight broods collected on different days, probably from different mothers; importantly, our results showed large variation between broods (Fig. 4F). For example, 1q ablation abolished organizer-dependent trunk ectoderm development in all embryos of one brood (n=12), while in another brood nearly all embryos (13/14) developed as headless but otherwise well-formed veligers. Such interbrood variation is likely to blame for Sweet’s failure to observe a defect in organizer induction. The between-brood variability may reflect genetic variation among wild snails, phenotypic plasticity in response to maternal or embryonic stress or simply stochastic variation in micromere signaling or 3D receptivity.

### 3D specification as an ultrasensitive response to a graded input from multiple cells

Our data support a model in which 3D specification depends on signaling from multiple micromeres, whose signaling activities could be simply additive, normally summing to surpass an activation threshold; 1q ablation may lower the signal level approximately to the threshold. We observed clear presence or absence of activated MAPK in 3D, with no evident intermediate states. This is consistent with an ultrasensitive (all or none) MAPK response to a threshold level of signal (Ferrell, 1996).

We are currently unable to judge the relative contributions of identified cells to 3D induction. In D quadrant isolates the combination of 1d and 2d appears necessary and sufficient to activate 3D MAPK. However, loss of these two cells in otherwise intact embryos appears to be compensated for by other cells. How many cells normally signal to 3D, and which ones? The occasional success of 3D specification in 1d-ablated D quadrant isolates weighs against a requirement for a minimum total number of micromeres higher than one quadrant’s worth. This fact, along with the variable results of 1q ablation in whole embryos, rules out a strict requirement for first-quartet cells *per se*: whatever the first quartet’s contribution to organizer induction, its absence can (sometimes) be compensated by the 2d cell, possibly with help from other cells.

### Evidence for separable inductive consequences of micromere signaling

As well as suggesting that 3D induction involves multiple intercellular signals, our results point to separable downstream effects. In contrast to the all-or-none 3D MAPK response, the lengthening of the 3D cell cycle observed in both D quadrant isolates and micromere ablations in whole embryos increases progressively with the removal of more cells. Unlike MAPK activation, cell cycle delay was observed in all D quadrant isolates, indicating that mitotic timing demands additional signaling beyond the threshold needed for MAPK activation. The distinct responses of MAPK activation and cell cycle control could be a clue to independent features of upstream signaling. Similarly, the response of MAPK activation in the micromeres showed a greater dependence on micromere signaling beyond the combination of (1d+2d), suggesting that this second wave of MAPK activation may reflect the operation of additional factors beyond the 3D MAPK activating signal.

### An unexpected role for 2d

The present study is the first to document a 3D-inducing role for any non-first-quartet cell. Juxtacrine signaling from 2d is certainly favored by its position (more accurately that of its daughters 2d^1^ and 2d^2^) with a large area of direct contact to 3D. However, daughters of 2a and 2c are also in direct (albeit more slim) contact with 3D (Sweet, 1996). Our results leave open the question as to whether the *I*.*obsoleta* 2d micromere has an intrinsically special property compared to other second-quartet micromeres; in other spiralian taxa with early D quadrant establishment, 2d is indeed intrinsically special (reviewed by Freeman and Lundelius, 1992; see also Lanza and Seaver, 2020), but such has not been shown in any gastropod. The contribution of 2d and/or other cells to 3D induction in *I. obsoleta* may not be unique: Boring (1989) reported a large minority of 1q-deleted embryos developing shell and/or foot structures in a cephalaspidean opisthobranch with equal cleavage. Future experiments should be done to identify all the sources of 3D-inducing signal in spiralian embryos.

### Not as evolved as we thought…

Polar lobe segregation into the D quadrant has apparently evolved in three molluscan lineages: in bivalves, in scaphopods, and in the hyperdiverse snail clade Caenogastropoda, including *Ilyanassa* and other neogastropods (Freeman and Lundelius, 1992). Experiments in two other caenogastropod taxa indicate surprising variation in polar lobe function. In *Bithynia*, PL removal abolished 3D organizer activity (Cather and Verdonk, 1974), while 1q ablation yielded complete trunks lacking heads (van Dam and Verdonk, 1982). In this case, PL inheritance indeed appears sufficient for autonomous 3D specification. In *Crepidula*, 1q ablation abolished 3D MAPK activation and organizer activity, while PL removal did not block organizer specification but rather randomized it to one 3Q cell or another (or sometimes more than one) (Henry et al., 2006; Henry et al., 2017). Further, ablation of the presumptive D macromere descended from the PL-inheriting quadrant founder cell in *Crepidula* resulted in functional replacement by another macromere (Henry et al., 2006). It seems that in different caenogastropod lineages, an ancestral mechanism conditionally specifying 3D has been supplemented or supplanted by segregated factors in vegetal cytoplasm that bias or override selection. The cell-autonomous choice may have taken over completely in *Bithynia*; in *Crepidula*, polar lobe material merely offers a strong but tentative suggestion. In *I. obsoleta*, the bias is absolute, but intercellular signaling is still required.

## MATERIALS AND METHODS

### Snails and Embryo Culture

*I. obsoleta* adults obtained from the Marine Resources Center (Marine Biological Laboratories, Woods Hole, MA) were kept in twenty-gallon tanks (ASW, Instant Ocean) (with an inch of aquarium rock substrate, sponge filtration and bubbler aeration) at room temperature (23°C (±1°C)) and fed frozen clam meat every other day. Embryos were collected from newly deposited egg capsules as described (Collier, 1981), and cultured at 23°C (±1°C) in groups of 3-15 in 35mm plastic petri dishes with 2-3ml of 0.2μm-filtered artificial sea water (FASW, Instant Ocean). For longer incubation, penicillin (100 units/ml) and streptomycin (200 μg/ml) were added, and embryos were moved to fresh FASW in a new dish every 2 days (Gharbiah et al. 2009).

### Microsurgery

For C or D quadrant isolations, the AB cell was killed by needle puncture during the early 2-cell stage, and its carcass removed; this was followed by killing and removing either the D cell or the C cell soon after completion of second cleavage. To isolate A or B quadrants, we killed the CD cell in the same manner at the 2-cell stage, and then killed one of the two indistinguishable remaining cells (A or B) shortly after their formation at second cleavage. For PL(−) quadrant isolations, polar lobes were removed at the trefoil stage (first cleavage) by gentle agitation in low-Ca^2+^/Mg^2+^ FASW (Collier, 1981). One cell at the 2-cell stage was killed and removed as described above, followed by another killing and removal after second cleavage. Micromere ablations were carried out during a short time window following the micromere quartet’s formation (10-25 minutes after apparently full constriction of cytokinetic furrows).

### Immunohistochemistry

MAPK activation was detected with a mouse monoclonal antibody (Sigma M8159) recognizing a diphosphorylated 11-amino-acid peptide in the conserved activation loop of ERK1/2, as previously described (Lambert and Nagy, 2001, Lambert and Nagy, 2003). Briefly, embryos were fixed in 3.7% formaldehyde in 90% FASW for 30-35 minutes. Embryos were dehydrated and stored in methanol at −20°C. Embryos were rehydrated by three 10-min washes in phosphate-buffered saline (PBS) + 0.1% Tween-20 (PBTw), blocked for at least 1 h in PBTw + 2% Bovine Serum Albumin (PBTw + BSA), then incubated 8–16 h at 4°C in primary antibody diluted in PBTw + 2% BSA. After primary incubation, embryos were washed 6 × 10 min in PBTw, then incubated with anti-mouse horseradish peroxidase (HRP)-linked secondary antibody for at least 2 h (23°C (±1°C)) with gentle agitation on a shaking platform, washed as above, and detected with Pierce HRP detection kit (Thermo Fisher) according to the manufacturer’s instructions. After HRP detection, embryos were incubated in DAPI (Molecular Probes;1 μg/ml) for 15 min (RT) then washed three times quickly in PBTw. PBTw was replaced with 80% glycerol: 1x PBS) for imaging.

### Imaging and analysis

In immunohistochemistry preparations the cell nuclei are visualized by DAPI staining. Larvae were reared at room temperature for seven days, anesthetized (Clement and Cather, 1957), and fixed using the same conditions as embryos. Larvae were scored with polarizing optics to visualize birefringent materials (shell, statocysts and operculum). Other larval structures (esophagus, stomodeum, intestine, retractor muscle, eye, velum) were identified according to the previous descriptions (Sweet, 1998; Atkinson, 1971). 3D division timing (i.e., the fourth micromere formation) was approximately determined by observing the embryos occasionally (every 15-20 minutes).

## ACKNOWLEDGEMENTS

We thank J. D. Lambert for valuable input on earlier versions of the manuscript. This work was supported by a National Science Foundation grant to L.M.N. (NSF IOS 0820564).

